# Determination of the absolute concentration of Rayleigh particles via scattering microscopy

**DOI:** 10.1101/2024.06.08.598040

**Authors:** Il-Buem Lee, Hyun-Min Moon, Jin-Sung Park, Se-Hwan Lee, Jaewon Lee, Sung Hun Park, Seungwoo Lee, Seok-Cheol Hong, Minhaeng Cho

## Abstract

Nanoparticles are crucial in diverse fields such as healthcare, electronics, and energy. Their small size allows them to cross biological barriers, enhancing drug delivery but also posing health risks. Accurate characterization of nanoparticles is essential for assessing their safety and efficacy, particularly in medical applications. Traditional methods such as dynamic light scattering and mass spectrometry have limitations in sensitivity and range of application. To address these challenges, we introduce the interferometric Concentration Analyzer and Ultrasmall Nanoparticle Tracker (iCAUNT), a technique for detecting and quantifying nanoparticles smaller than one-tenth of the imaging light wavelength. As a non-invasive method, iCAUNT provides precise size and concentration measurements of biological and synthetic nanoparticles, offering significant potential for diagnostics, therapeutics, and broader nanoscience applications.

## Main Text

Nanoparticles (NPs) have emerged as critical elements across a wide range of industries, including healthcare and medicine, electronics, energy, food and agriculture, and cosmetics. In healthcare and medicine, NPs are revolutionizing biomedical imaging, cancer therapy, diagnostic assays, and vaccine development. Many functional elements of biological systems, such as protein complexes, lipid droplets, intracellular cargos, exosomes, and viral particles, are natural biological NPs. The small size of synthetic NPs enables them to easily cross biological barriers, making them useful for applications like drug delivery. However, this also poses potential risks for adverse effects on human health and the environment. Therefore, it is essential to characterize NPs to assess their physical properties, toxicity, and long-term health effects, thereby defining safe levels of human exposure (1, 2).

Determination of the size and concentration of NPs is crucial in environmental and therapeutic contexts due to their potential biological impacts and the dose-dependent nature of their interactions with biological systems. Accurate quantification of NP concentration is vital for dosing in medical applications such as targeted drug delivery, where the optimal dosage is critical for achieving therapeutic effectiveness while minimizing toxicity. Additionally, NPs at high concentrations, even if benign individually, may aggregate and cause serious health effects such as inflammation or cytotoxicity (3). Understanding and controlling NP concentration is therefore essential for minimizing risks associated with overexposure and ensuring environmental safety. This highlights the need for robust methodologies to reliably assess and monitor NP concentrations across various settings, guaranteeing effective clinical outcomes and adherence to environmental standards.

Traditional methods for characterizing and enriching NPs aim to provide insights into their physical, biochemical, and biological properties (4). Techniques such as mass spectrometry and gel electrophoresis are used to analyze molecular composition and purity of NPs, while biological assays like surface plasmon resonance (SPR) and enzyme-linked immunosorbent assay (ELISA) focus on their interactions with antibodies. The distribution of particle size is examined using dynamic light scattering (DLS) and analytical ultracentrifugation, whereas advanced non-optical microscopy offers high-resolution imaging of NP morphology.

Despite their utility, these methods have limitations: high-resolution imaging requires complex sample preparation, DLS may struggle with polydisperse samples, and biological assays might not accurately predict *in vivo* immune responses. Additionally, variability in nano-material production from batch to batch and distinct sample preparation requirements for different methods can affect the scalability and reproducibility of measurements. These challenges highlight ongoing issues in the field of NP research (5).

Recently, interferometric scattering microscopy (iSCAT) has attracted significant attention as a sensitive, high-speed, and long-term imaging tool for label-free detection (fig. S1) (6, 7). iSCAT can detect both dielectric and metallic NPs with a lower size limit for gold NPs to a few nanometers and proteins down to tens of kilodaltons or lower by utilizing transient binding events to the solution-glass interface, providing unprecedented resolution and sensitivity in size measurement (6, 8-11). Label-free detection by iSCAT enables the observation of the entire population of biological NPs in their native state, without the artifacts and photo-toxicity associated with fluorescence labeling, and independent of the presence or quality of specific labels (12, 13). The high sensitivity of iSCAT also expands its dynamic range in time domain significantly. As a result, iSCAT excels in delivering detailed insights into nanoscale biological processes, such as receptor-ligand interactions and endocytosis, by tracking these events with high temporal resolution (7, 14-16).

More recently, Nanofluidic Scattering Microscopy (NSM) has emerged as a powerful tool for real-time, label-free imaging and precise determination of the size, mass, and hydrodynamic properties of single diffusing biomolecules and NPs within nanofluidic channels. By reducing the rapid diffusion of proteins, NSM enhances observation capabilities, enabling detailed study of these entities (17). The integration of iSCAT microscopy with nanoparticle tracking analysis (NTA), named iNTA, has enabled size determination down to 10 nm for metallic NPs and 40 nm for dielectric NPs in suspension, as well as refractive index measurement for liposomes (18).

Holographic NTA, leveraging a *k*-space holographic sensing platform, has extended the observation volume and allowed for 3D single-particle tracking (SPT) (19). Rytov microscopy for nanoparticles identification (RYMINI), utilizing a modified Hartmann mask, has enabled label-free imaging of NPs with more than 90% accuracy in classification (20).

These advancements have promoted label-free detection of NPs in solution, as detailed in table S1. However, challenges remain: the lower bound of detectable size is still limited and the absolute concentration measurements of NPs have not been achieved partly because of limited tracking range, undefined imaging depth, and incomplete sensing of NPs within the imaging volume.

In this work, we focused on detection of dielectric NPs smaller than one-tenth of the wavelength of imaging light and aimed to accurately measure the diffusion coefficient and effective imaging volume through SPT analysis. By combining these approaches, we accurately measured the size of small (< 40 nm) NPs that scatter light strictly in the Rayleigh regime, referred to as Rayleigh particles (RPs), and determined the absolute concentrations of RPs, including those composed of biological material. This all-optical, non-invasive approach enables quantitative characterization of delicate biological complexes in natural aqueous environments, a significant advancement.

We have named our technique the interferometric Concentration Analyzer and Ultrasmall Nanoparticle Tracker (iCAUNT). We envisage that iCAUNT would have substantial applications in biomolecular diagnostics and therapeutics, as well as in general nanoscience and nanotechnology, by facilitating high-throughput, label-free characterization of biological Rayleigh particles (bRPs).

### Label-free imaging of Rayleigh particles in suspension

As noted, iSCAT microscopy is a highly sensitive imaging technique that enables shot-noise-limited, high-speed detection of NPs as diffraction-limited spots (Fig. 1) (6, 8). An array of contrasting and filtering mechanisms (Fig. 1B and fig. S2), along with differential attenuation of output fields, reference and scattering, for optimal signal acquisition (partial mask (PM) in fig. S1), have made iSCAT even more sensitive, allowing for the sensing of RPs (fig. S3) (9, 10). In this work, we have made further improvements to detect RPs in suspension, identify them more reliably, and elicit detailed physical information about them.

**Fig. 1.**
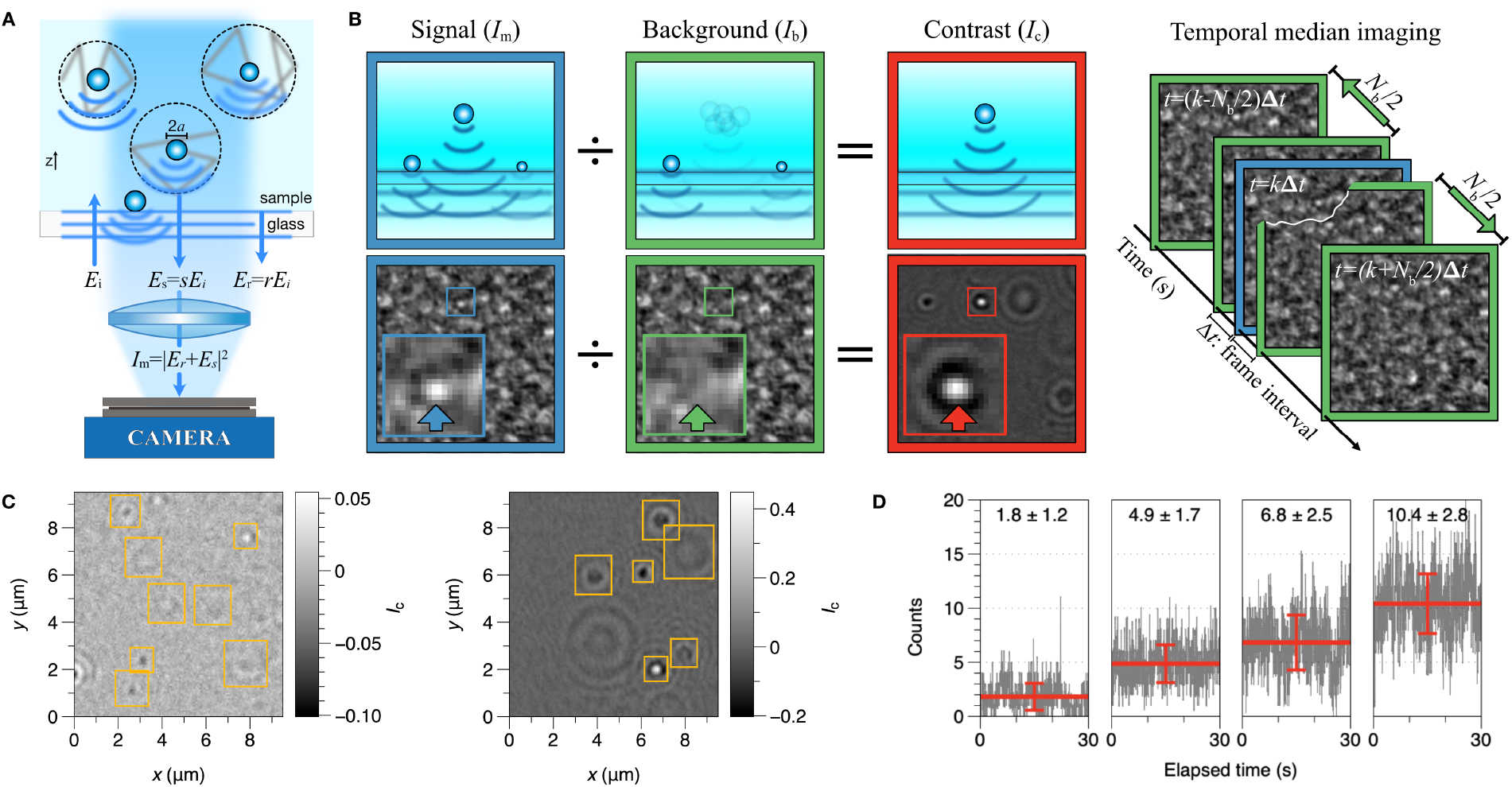
Label-free imaging for nanoparticles and demonstration of single particle detection. (A) Schematic illustration of iSCAT detection of mobile NPs dispersed in solution. The side view shows the interferometric measurement of reference (*E*_r_) field from the glass surface and scattering field (*E*_s_) from NPs, encircled in dashed line to delineate the diffusion length. (B) Illustration of the scheme to obtain a processed image (*I*_c_) by removing the static background (*I*_b_) from a measured image (*I*_m_) via temporal median filtering (pixel-to-pixel division). The static background image was created by averaging *N*_b_ consecutive frames symmetrically selected before and after the *one-frame* image to be processed (inset: zoomed-in images of the boxed area with the arrows pointing to the location of an NP). (C) Representative iSCAT contrast images of PS20 (left) and PS40 (right) beads with the grayscale legend representing iSCAT contrast and the yellow boxes marking the particles identified via our YOLOv4-based method. (D) Number counts for PS20 in suspension at different concentrations ([PS20] = 25, 50, 100, and 200 pM from left to right). The count of PS20 fluctuates in time (gray) with the mean value and error shown in red. See also figs. S1 to S8 and movies S1 to S5.

First, we systematically adjusted the parameters of image processing and analysis. One of the signal processing strategies frequently used in iSCAT to capture weak signals is the time differential method, which effectively removes stationary or slow-varying backgrounds and highlights dynamic, fast-changing features (21). When detecting mobile particles in suspension, it is preferable to obtain a stationary background image, which is used to eliminate non-uniform static backgrounds from measured images taken at different time points and to reveal NPs as distinct spots. Despite motion-induced blurring, fast-moving particles in suspension offer several detection advantages: by focusing on particles floating a few μm away from the glass surface, inhomogeneities originating from the glass surface are largely absent; non-uniformity due to scattering from nearby dynamic particles can be effectively eliminated by using a background image averaged over an optimally selected time period (movies S1 to S3). The optimal number (*N*_b_) of frames to average for minimal background noise can be determined experimentally for NPs of different size (figs. S4 and S5 and movie S4). For example, the *N*_b_ values for PS20, PS30, and PS40 are 20, 100, and 320, respectively, with the ratios *N*_b_(PS30)/*N*_b_(PS20) = 5 and *N*_b_(PS40)/*N*_b_(PS20) = 16, which can be explained as the quartic power of the ratio of particle size (Supplementary Text). This background preparation method allowed us to capture particles clearly by sufficiently smearing out scattering noise from dynamic particles.

Second, we theoretically created the point spread function (PSF) of an NP of each kind at different focal positions, which reproduces the corresponding experimental observations (22-24). Using these data sets, we generated a series of videos that simulate the motion of a group of NPs in suspension, adding a certain level of artificial noise and allowing spatial overlap of their images. These videos, along with the known information about the NPs, served as the ground truth for training our YOLOv4-based deep learning algorithm, which we used to identify NPs and detect their 3D positions from noisy video data (25). Since a high NA objective was used in our setup to collect as many photons as possible, the depth of field was limited to a few microns at most. This caused concentric sharp fringes to evolve rapidly as particles moved away from the focal plane, which proved advantageous for particle identification (due to the richness of information) and particle counting (due to shallow sectioning; see below) (26, 27). It is challenging to identify an NP from its highly defocused image with conventional intensity-threshold-based methods (28), but our deep learning-based approach overcomes this challenge, as demonstrated (Fig. 1C and fig. S6). Our PSF-based particle detection algorithm, which utilizes Mie scattering theory to model a wide range of particles and employs YOLOv4, is well-suited for tracking multiple NPs simultaneously in movie clips generated by iSCAT imaging (fig. S7). This capability is particularly important for measuring the concentration of NPs in suspension (Fig. 1D and movie S5). Representative time traces and frequentness of number counts of all RPs tested in this study are shown at different concentrations in fig. S8.

Third, we accurately characterized the behavior of NPs in Brownian motion. To determine the absolute concentration of RPs in suspension, we determined the depth of field for particles of different size. While tracking the vertical diffusion of an NP is a matter of subtlety and ambiguity, lateral diffusion can be precisely and directly characterized via imaging. We observed that the lateral and vertical dimensions are equivalent, meaning the diffusion coefficient *D* measured from lateral diffusion should be the same for vertical diffusion (28-30). Using a large set of particle trajectories, we reliably measured the depth of field. Moreover, we determined diffusion coefficient *D* using the covariance-based estimator (CVE) instead of the mean square displacement (MSD) analysis, as the former is unbiased and more reliable (31, 32). With these advancements, we effectively tackled the problems of tracking RPs and accurately gauging their concentrations.

### Hydrodynamic characterization of Rayleigh particles by SPT analysis

The iSCAT-based SPT method, enhanced with the aforementioned developments, permits precise and reliable measurement of the size of RPs in suspension. The CVE is an unbiased estimator of *D* and does not require a long particle trajectory, unlike the MSD-based approach, making it versatile and robust under various experimental conditions (31, 32). Thus, using the Stokes-Einstein relation, the radius of a diffusing particle can be determined more reliably if the viscosity of the surrounding medium is known, or the nanoscale viscosity can be measured if NPs of precise size are available. The characteristics of SPT for various RPs are graphically summarized in fig. S9.

To ensure the precision of our SPT analysis, we used National Institute of Standards and Technology (NIST)-certified polystyrene (PS) beads for their size uniformity and accuracy. We measured the diffusion coefficient of PS beads suspended in pure water and glycerol-water mixtures (Fig. 2A and movies S6 to S9). Our findings indicate that particle mobility in solution is inversely correlated with particle size and the viscosity of the solution. We also measured the viscosity of a solution using the formula *η* = (*D*_0_/*D*) *η*_w_ (*η*_w_: viscosity of pure water) with high-precision NIST PS beads. The SPT-based nano-viscometry yielded viscosity values of 6.2, 10.4, and 19.2 cP for 50%, 60%, and 70% glycerol, respectively (Fig. 2A, fig. S10, and movies S7 to S9), which are in good agreement with theoretical values (33). We found that the distribution of particle sizes, derived from measured diffusion coefficients and calibrated solution viscosity, aligned well with the average particle size reported by the manufacturer (Fig. 2B and fig. S10).

**Fig. 2.**
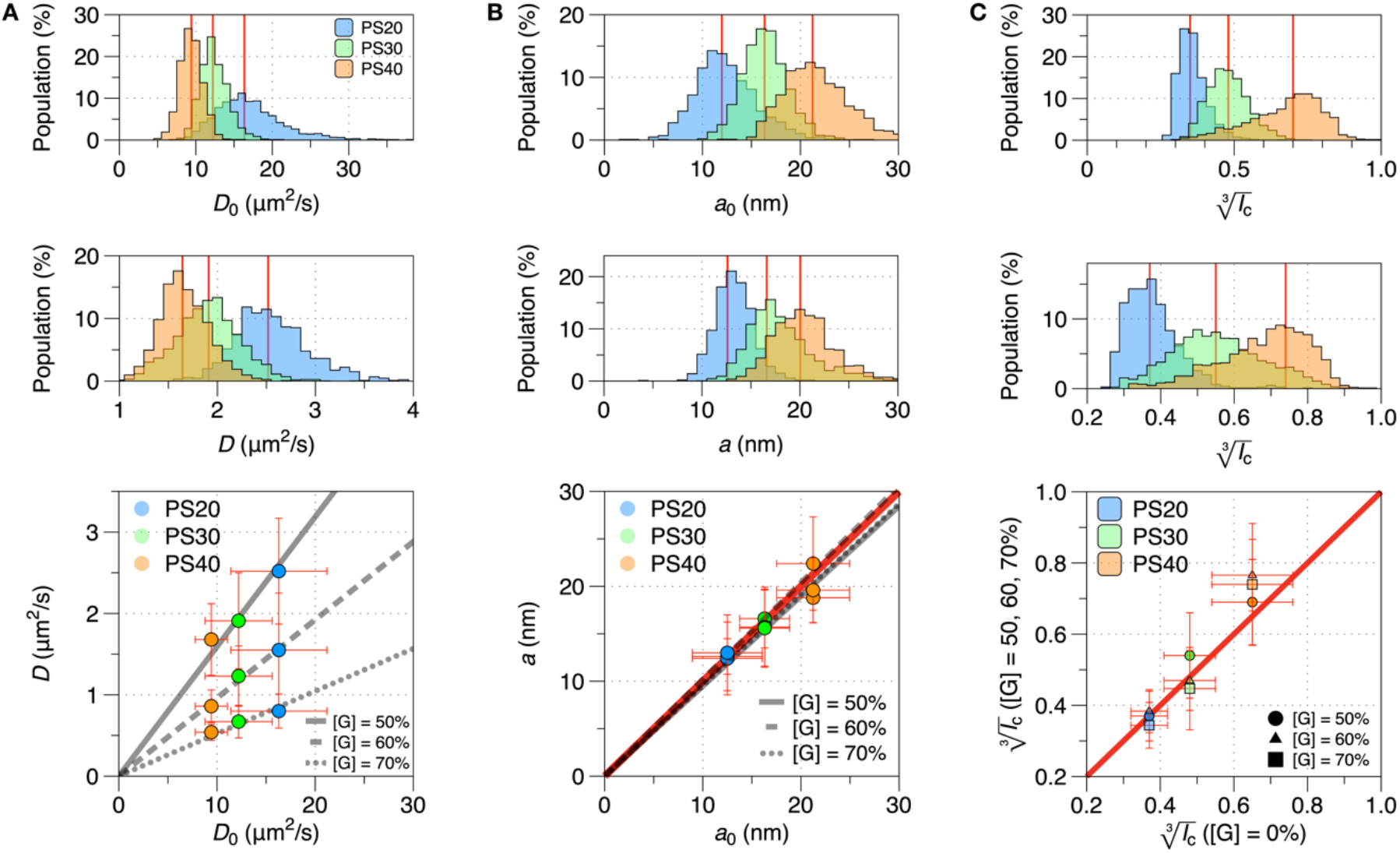
SPT analysis for the standard PS beads in water and 1:1 glycerol-water mixture. (A) Distribution of the diffusion coefficient of RPs in pure water (*η*_w_ ∼ 1 cP, top) and glycerol-water mixture (50% glycerol (v/v): [G] = 50%, middle). The median values of diffusion coefficients of PS beads (*D*_0_ in water, *D* in [G] = 50%) are (16.2, 2.51) for PS20, (12.2, 1.91) for PS30, and (9.4, 1.64) for PS40, in μm^2^/s, respectively. Scatter plot (bottom) of (*D*_0_, *D*) highlighting the effects of RP size and glycerol concentration on RP mobility. The inverse of the slope of each line fit indicates the viscosity of the corresponding solution (*η* = (*D*_0_/*D*)*η*_w_): *η*([G] = 50%) = 6.2 cP (solid), *η*([G] = 60%) = 10.4 cP (dashed), and *η*([G] = 70%) = 19.2 cP (dotted). (B) Distribution of the size of RP in pure water (top) and glycerol-water mixture ([G] = 50%, middle). The median sizes of PS beads (*a*_0_ in water, *a* in [G] = 50%) are (12.5, 12.6) for PS20, (16.3, 16.6) for PS30, and (21.2, 19.3) for PS40, in unit of nm, respectively. Scatter plot (bottom) of (*a*_0_, *a*) showing quantitative agreement in size measurement, as depicted by a line with a slope of 1 (red). (C) Distribution of the cube root of positive contrast from RP measured in pure water (top) and glycerol-water mixture ([G] = 50%, middle). The median contrasts 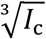 in water, 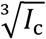 at [G] = 50%) are (0.36, 0.37) for PS20, (0.48, 0.55) for PS30, and (0.70, 0.71) for PS40, respectively. Scatter plot of measured cube-root contrasts 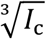 in water, 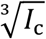 at [G] = 50, 60, 70%), displaying the correlation of unit linearity as shown in red. The red solid lines drawn on the distributions of the diffusion coefficient (*D*_0_, *D*), particle size (*a*_0_, *a*), and contrast 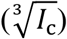 indicate the median values of the respective quantities. See also figs. S9 and S10 and movies S6 to S9.

We also acquired the distribution of iSCAT contrast for PS beads of different size in solutions with various viscosity levels in order to assess the consistency of measured iSCAT contrasts. We found that the iSCAT contrasts of these RPs were not considerably altered by the content of glycerol, and thus the refractive index of the medium (Fig. 2C and fig. S10). Notably, our iSCAT microscopy succeeded in detecting and tracking PS beads of diameter = 20 nm in suspension, an achievement not previously accomplished by NTA techniques or other iSCAT works to date (18, 19, 34).

### Characterization of biological Rayleigh particles

We also demonstrated that our iSCAT-based SPT assay could characterize bRPs. bRPs are diverse and heterogeneous in substance and non-uniform and complex in composition, including simple proteins, lipid droplets, virus particles, and artificial composites such as DNA origami. Various physiological conditions such as acidity, ionic strength, and molecular crowding must be controlled for their stability. Glycerol is a convenient agent to add to protein samples to adjust the degree of molecular crowding. Thus, we examined the dynamics of bRPs diffusing in concentrated glycerol solutions (35).

As expected, we found that the iSCAT contrast of bRPs increased and their mobility decreased with particle size. The mean radii deduced by tracking measurement were 10.7 ± 4.7 nm for IgM, 20.1 ± 8.2 nm for SV40, and 23.5 ± 7.3 nm for DNA origami (Fig. 3A and movie S10). These values are in good agreement with those obtained by conventional measurements (36-39). We also observed that scattering by bRPs was much smaller than by PS beads; for instance, the iSCAT contrast for SV40 was half that of PS40, despite their similar sizes (Fig. 3B). This indicates that what matters in scattering is not a mere size of scatterer but the combination of scatterer size and relative refractive index, as shown in Rayleigh scattering formula (40, 41).

**Fig. 3.**
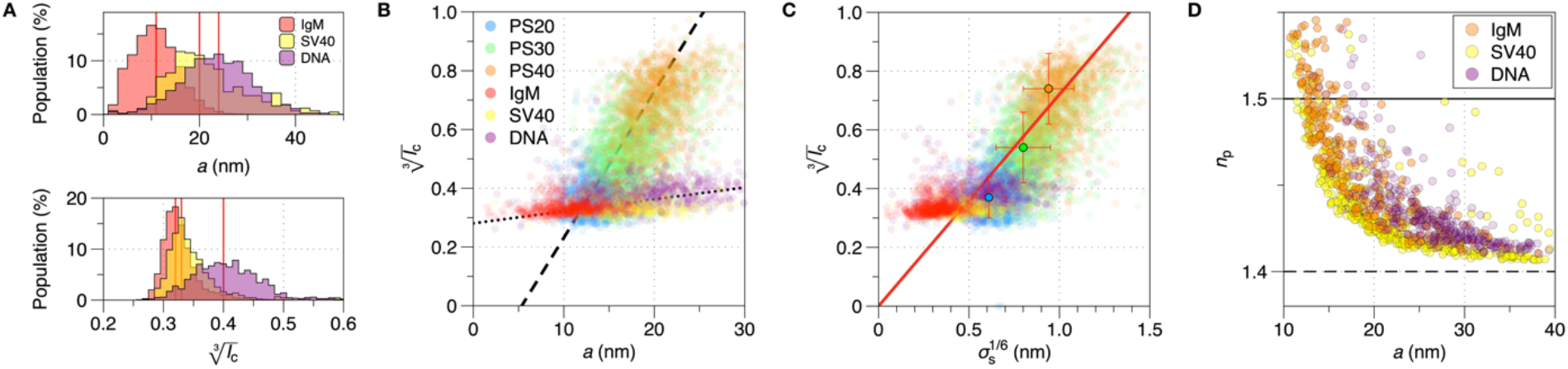
Characterization of biological Rayleigh particles. (A) Distributions of the measured size (top) and cube-root contrast (bottom) of immunoglobulin M (IgM), simian virus 40 (SV40), and DNA origami in glycerol-water mixture ([G] = 50%). The median values of contrast and radius in unit of nm, 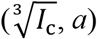, are (0.33, 10.7) for IgM, (0.34, 20.1) for SV40, and (0.40, 23.5) for DNA origami, respectively. The red solid lines indicate the median values in each distribution. (B) Scatter plot of 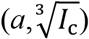 for all RP samples tested with trend lines for PS beads (dashed) and for biological RPs (dotted), indicating that scattering strength correlates with particle size and the slope of correlation differs distinctively for solid PS beads and porous bRPs. (C) Scatter plot of 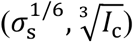 for all RP samples tested with the universal trend line (mean 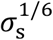 of PS beads: 0.61 ± 0.09 nm for PS20 (blue dot), 0.80 ± 0.15 nm for PS30 (green dot), and 0.94 ± 0.14 nm for PS40 (orange dot)), which is given by 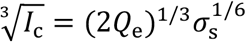 with *Q*_e_ = 0.18 /nm. The 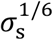 of bRP is similar to that of PS20, indicating that the refractive index as well as particle size is important for scattering field. (D) Scatter plot of the measured refractive index and size of bRP (*a, n*_p_). The dashed and solid lines indicate the values of the refractive index of [G] = 50% and typical protein material, respectively (42-44). See also fig. S9 and movie S10.

The iSCAT contrast is related to the scattering cross section *σ*_s_ by 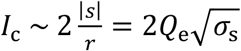 where *Q*_e_ is the detection efficiency, which depends on the setup and is common for all our measurements. Using the data from the standard PS beads with a well-defined size and refractive index, we obtained *Q*_e_ ∼ 0.18 /nm (6, 18). Using this value of *Q*_e_, the measured *I*_c_, and the radius of bRP (*a*), we could determine the refractive index of bRP, which varies from 1.4 to 1.5, depending on *a* (Fig. 3, C and D; see also Supplementary Text). The measured refractive indices are in good agreement with typical values for various biological samples (42-44). Here, it is worth mentioning that iSCAT allows us to assess the optical properties of NPs in addition to their physical size with high precision, providing critical information for a broad range of scientific and technological applications.

### Determination of the absolute concentration of Rayleigh particles

It is a great challenge to count Rayleigh particles precisely and, more so, to determine their absolute concentrations in general. A noise-suppressed holographic image of an RP is a prerequisite for reliable analysis (1st and 2nd advances above). Accurate trajectory tracking of RPs for an extended period and objective analysis of their diffusion provide the information needed to determine particle concentration (3rd advance above).

Small RPs (e.g., PS20) scatter light weakly and move rapidly, making the focal region that yields a detectable signal very thin. This region holds fewer RPs for the same concentration and is traversed by small RPs too transiently. Thus, boosted sensitivity becomes pivotal in this regime for rapid data acquisition, and the CVE-based determination of *D* is particularly useful for the short trajectories exhibited by small particles. Conversely, larger RPs, like PS40, move slowly and scatter light strongly, making measurements easier. However, they may be detectable at greater depths beyond the imaging volume, contributing to dynamic noise (Fig. 1). To create a quasi-static background from images with many dynamic particles, we accumulated numerous image frames (*N*_b_) in which dynamic particles migrate over many pixels, causing the scattering signal detected on each pixel to be smeared below the typical measurement noise in each pixel.

We measured the diffusion length, which quantifies the average distance a particle travels due to Brownian motion over time. The observation time can be converted to the range of particle’s vertical motion, serving as a proxy for the effective imaging depth, *l*_z_ (45). The *l*_z_ values measured were 1.45 ± 0.18 μm for PS20, 3.07 ± 0.26 μm for PS30, and 4.09 ± 0.24 μm for PS40 (Fig. 4A). We also measured *l*_z_ for bRPs: 0.31 ± 0.03 μm for IgM, 0.29 ± 0.05 μm for SV40, and 0.97 ± 0.15 μm for DNA origami (fig. S11 and table S2). We also acquired the distributions of *peak* iSCAT contrasts 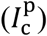 for all RPs tested (Fig. 4B). The interquartile ranges (IQR) of the negative and positive lobes of peak iSCAT contrast (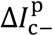and 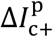) progressively increased with nominal iSCAT contrast, more significantly in the positive lobe.

**Fig. 4.**
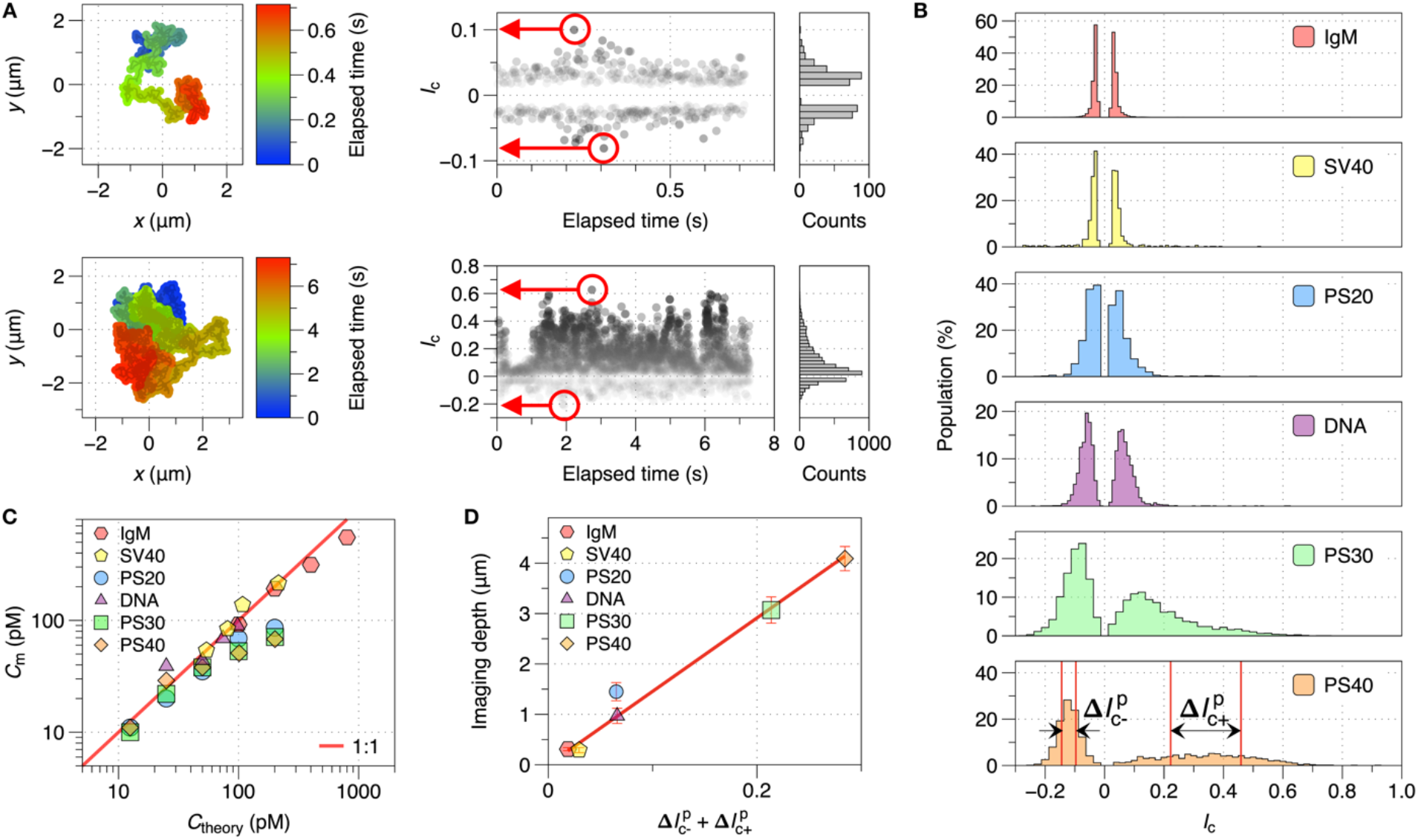
Absolute concentrations of Rayleigh particles determined by SPT and imaging volume measurement. (A) Representative particle trajectories in 2D (left) and time traces of iSCAT contrast (right) from PS beads (PS20: top; PS40: bottom) in suspension. The larger the size of a particle is, the slower and longer it travels and the stronger the iSCAT contrast is. The largest positive and negative contrast values in each time trace were chosen as the representative values of the trace (red arrows). (B) Distributions of the *peak* contrasts 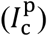 for all RPs tested. 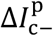 and 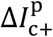 represent the IQR of the negative and positive lobes of *peak* iSCAT contrast distributions: 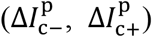 are (0.009, 0.010) for IgM, (0.014, 0.015) for SV40, (0.028, 0.037) for PS20, (0.029, 0.037) for DNA origami, (0.047, 0.157) for PS30, and (0.048, 0.237) for PS40, respectively. (C) Scatter plot of the supposed and measured absolute concentrations (*C*_theory_, *C*_m_) for all RPs tested, showing excellent agreement between the two concentrations for all RPs tested. The same plots on a linear y-scale, with one plot per sample, are shown in fig. S12, including error bars representing the standard deviation of each measured concentration. (D) Scatter plot of the ‘full width’ of peak contrast distribution vs. effective imaging depth 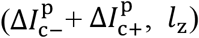. The linear relation indicates that particles with stronger scattering and thus scattering strength are detected over a larger vertical range yielding a thicker imaging volume. See also figs. S11 and S12 and tables S2 and S3.

The absolute concentration of each sample was obtained by dividing the average counts of RPs by the effective imaging volume, which was determined by *l*_z_. Intriguingly, the absolute concentrations measured in this way showed excellent quantitative agreement with the values reported by the manufacturer. This indicates that we have successfully demonstrated the first scattering-based label-free measurement technique for determining the absolute concentration of NPs smaller than 50 nm in diameter, a range of size where popular commercial methods such as the NTA technique fail to provide reliable counts (Fig. 4C and fig. S12).

Curiously, the effective imaging depth is linearly correlated with the ‘full width’ (sum of the IQRs of the positive and negative lobes) of the peak contrast distribution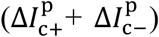. This relationship indicates that the stronger the scatterer, the greater the imaging depth and the broader the peak value of iSCAT contrast would be. This correlation makes sense because a strong scatterer can be identified farther away from the focal plane, resulting in a thicker focal volume, and the peak value of iSCAT contrast in each trajectory would be widely distributed due to the variability in the paths taken by the scatterer − some passing directly through the focal plane and others merely skimming the edge of the imaging space (46). By obtaining this correlation parameter, we could determine the imaging depth just from the distribution of iSCAT contrast, providing a convenient proxy for the effective imaging depth (Fig. 4D and table S3).

## Conclusion

This study underscores the straightforwardness, sensitivity, quantitativeness, and versatility of the iCAUNT technique in tracking and characterizing NP dynamics in biological samples. This technique enables real-time, label-free visualization, allowing for high-precision quantitative analysis of NP size, concentration, and refractive index. The incorporation of deep-learning-based analysis for improved accuracy in particle detection and tracking further enhances the current technology via holographic imaging. The nanoscopic method presented here is valuable for investigating the dynamics of small NPs and their interactions in complex biological environments, providing key insights into biological phenomena and the development of advanced diagnostic and therapeutic tools. The ability to examine viral and other biological particles without labeling is poised to transform pathogen detection and analysis, facilitating rapid and precise diagnostics. We envisage that the analysis of NPs in body fluid using this technology will lead to much-anticipated breakthroughs in biological science and medicine.

## Author contributions

Conceptualization: SCH, MC, Investigation: IBL, Methodology: IBL and HMM, Analysis software: IBL, HMM, Deep-learning software: SHL, Validation: HMM, Resources: JSP, DNA origami design, synthesis, and characterization: JL, SHP, SL, Writing-original draft: IBL, Writing-review & editing: IBL, JSP, SCH, MC, Supervision: SCH, MC

## Data and materials availability

All data necessary to support the conclusions are available in the manuscript or supplementary materials.

## References and Notes

1. A. C. Anselmo, S. Mitragotri, Nanoparticles in the clinic. Bioeng. Transl. Med. 1, 10–29 (2016).

2. P. H. Hoet, I. Brüske-Hohlfeld, O. V. Salata, Nanoparticles – known and unknown health risks. J. Nanobiotechnology 2, 12 (2004).

3. W. H. D. Jong, P. J. Borm, Drug delivery and nanoparticles: applications and hazards. Int. J. Nanomed. 3, 133–149 (2008).

4. S. Nooraei, H. Bahrulolum, Z. S. Hoseini, C. Katalani, A. Hajizade, A. J. Easton, G. Ahmadian, Virus-like particles: preparation, immunogenicity and their roles as nanovaccines and drug nanocarriers. J. Nanobiotechnology 19, 59 (2021).

5. S. Mourdikoudis, R. M. Pallares, N. T. K. Thanh, Characterization techniques for nanoparticles: comparison and complementarity upon studying nanoparticle properties. Nanoscale 10, 12871–12934 (2018).

6. K. Lindfors, T. Kalkbrenner, P. Stoller, V. Sandoghdar, Detection and Spectroscopy of Gold Nanoparticles Using Supercontinuum White Light Confocal Microscopy. Phys. Rev. Lett. 93, 037401 (2004).

7. P. Kukura, H. Ewers, C. Müller, A. Renn, A. Helenius, V. Sandoghdar, High-speed nanoscopic tracking of the position and orientation of a single virus. Nat. Methods 6, 923–927 (2009).

8. M. Piliarik, V. Sandoghdar, Direct optical sensing of single unlabelled proteins and super-resolution imaging of their binding sites. Nat. Commun. 5, 4495 (2014).

9. D. Cole, G. Young, A. Weigel, A. Sebesta, P. Kukura, Label-Free Single-Molecule Imaging with Numerical-Aperture-Shaped Interferometric Scattering Microscopy. ACS Photonics 4 (2017).

10. G. Young, N. Hundt, D. Cole, A. Fineberg, J. Andrecka, A. Tyler, A. Olerinyova, A. Ansari, E. G. Marklund, M. P. Collier, S. A. Chandler, O. Tkachenko, J. Allen, M. Crispin, N. Billington, Y. Takagi, J. R. Sellers, C. Eichmann, P. Selenko, L. Frey, R. Riek, M. R. Galpin, W. B. Struwe, J. L. P. Benesch, P. Kukura, Quantitative mass imaging of single biological macromolecules. Science 360, 423–427 (2018).

11. M. Dahmardeh, H. M. Dastjerdi, H. Mazal, H. Köstler, V. Sandoghdar, Self-supervised machine learning pushes the sensitivity limit in label-free detection of single proteins below 10 kDa. Nat. Methods 20, 442–447 (2023).

12. Y. Li, W. B. Struwe, P. Kukura, Single molecule mass photometry of nucleic acids. Nucleic Acids Res. 48, gkaa632 (2020).

13. F. Soltermann, E. D. B. Foley, V. Pagnoni, M. Galpin, J. L. P. Benesch, P. Kukura, W. B. Struwe, Quantifying Protein–Protein Interactions by Molecular Counting with Mass Photometry. Angew. Chem. Int. Ed 59, 10774–10779 (2020).

14. Y.-H. Lin, W.-L. Chang, C.-L. Hsieh, Shot-noise limited localization of single 20 nm gold particles with nanometer spatial precision within microseconds. Opt. Express 22, 9159 (2014).

15. Y.-F. Huang, G.-Y. Zhuo, C.-Y. Chou, C.-H. Lin, W. Chang, C.-L. Hsieh, Coherent Brightfield Microscopy Provides the Spatiotemporal Resolution To Study Early Stage Viral Infection in Live Cells. ACS Nano 11, 2575–2585 (2017).

16. R. W. Taylor, R. G. Mahmoodabadi, V. Rauschenberger, A. Giessl, A. Schambony, V. Sandoghdar, Interferometric scattering microscopy reveals microsecond nanoscopic protein motion on a live cell membrane. Nat. Photonics 13, 480–487 (2019).

17. B. Špačková, H. K. Moberg, J. Fritzsche, J. Tenghamn, G. Sjösten, H. Šípová-Jungová, D. Albinsson, Q. Lubart, D. van Leeuwen, F. Westerlund, D. Midtvedt, E. K. Esbjörner, M. Käll, G. Volpe, C. Langhammer, Label-free nanofluidic scattering microscopy of size and mass of single diffusing molecules and nanoparticles. Nat. Methods 19, 751–758 (2022).

18. A. D. Kashkanova, M. Blessing, A. Gemeinhardt, D. Soulat, V. Sandoghdar, Precision size and refractive index analysis of weakly scattering nanoparticles in polydispersions. Nat. Methods 19, 586–593 (2022).

19. U. Ortiz-Orruño, R. Quidant, N. F. van Hulst, M. Liebel, J. O. Arroyo, Simultaneous Sizing and Refractive Index Analysis of Heterogeneous Nanoparticle Suspensions. ACS Nano 17, 221–229 (2023).

20. M. Nguyen, P. Bonnaud, R. Dibsy, G. Maucort, S. Lyonnais, D. Muriaux, P. Bon, Label-Free Single Nanoparticle Identification and Characterization in Demanding Environment, Including Infectious Emergent Virus. Small, e2304564 (2023).

21. J. O. Arroyo, D. Cole, P. Kukura, Interferometric scattering microscopy and its combination with single-molecule fluorescence imaging. Nat. Protoc. 11, 617–633 (2016).

22. R. G. Mahmoodabadi, R. W. Taylor, M. Kaller, S. Spindler, M. Mazaheri, K. Kasaian, V. Sandoghdar, Point spread function in interferometric scattering microscopy (iSCAT) Part I: aberrations in defocusing and axial localization. Opt. Express 28, 25969 (2020).

23. J. Dong, D. Maestre, C. Conrad-Billroth, T. Juffmann, Fundamental bounds on the precision of iSCAT, COBRI and dark-field microscopy for 3D localization and mass photometry. J Phys. D. Appl. Phys. 54, 394002 (2021).

24. K. Žambochová, I.-B. Lee, J.-S. Park, S.-C. Hong, M. Cho, Axial profiling of interferometric scattering enables an accurate determination of nanoparticle size. Opt. Express 31, 10101 (2023).

25. A. Bochkovskiy, C.-Y. Wang, H.-Y. M. Liao, YOLOv4: Optimal Speed and Accuracy of Object Detection. arXiv, doi: 10.48550/arxiv.2004.10934 (2020).

26. S. F. Gibson, F. Lanni, Experimental test of an analytical model of aberration in an oil-immersion objective lens used in three-dimensional light microscopy. J. Opt. Soc. Am. 8, 1601 (1991).

27. P. Török, P. Varga, Electromagnetic diffraction of light focused through a stratified medium. Appl. Optics 36, 2305 (1997).

28. K. A. Rose, M. Molaei, M. J. Boyle, D. Lee, J. C. Crocker, R. J. Composto, Particle tracking of nanoparticles in soft matter. J. Appl. Phys. 127, 191101 (2020).

29. H. Qian, M. P. Sheetz, E. L. Elson, Single particle tracking. Analysis of diffusion and flow in two-dimensional systems. Biophys. J. 60, 910–921 (1991).

30. X. Michalet, Mean square displacement analysis of single-particle trajectories with localization error: Brownian motion in an isotropic medium. Phys. Rev. E 82, 041914 (2010).

31. C. L. Vestergaard, P. C. Blainey, H. Flyvbjerg, Optimal estimation of diffusion coefficients from single-particle trajectories. Phys. Rev. E 89, 022726 (2014).

32. C. L. Vestergaard, Optimizing experimental parameters for tracking of diffusing particles. Phys. Rev. E 94, 022401 (2016).

33. J. B. Segur, H. E. Oberstar, Viscosity of Glycerol and Its Aqueous Solutions. Ind. Eng. Chem. 43, 2117–2120 (1951).

34. J. Gross, S. Sayle, A. R. Karow, U. Bakowsky, P. Garidel, Nanoparticle tracking analysis of particle size and concentration detection in suspensions of polymer and protein samples: Influence of experimental and data evaluation parameters. Eur. J. Pharm. Biopharm. 104, 30–41 (2016).

35. V. Vagenende, M. G. S. Yap, B. L. Trout, Mechanisms of Protein Stabilization and Prevention of Protein Aggregation by Glycerol. Biochemistry 48, 11084–11096 (2009).

36. J. K. Armstrong, R. B. Wenby, H. J. Meiselman, T. C. Fisher, The Hydrodynamic Radii of Macromolecules and Their Effect on Red Blood Cell Aggregation. Biophys. J. 87, 4259– 4270 (2004).

37. R. R. Akhouri, S. Goel, H. Furusho, U. Skoglund, M. Wahlgren, Architecture of Human IgM in Complex with P. falciparum Erythrocyte Membrane Protein 1. Cell Rep. 14, 723–736 (2016).

38. M. G. M. van Rosmalen, C. Li, A. Zlotnick, G. J. L. Wuite, W. H. Roos, Effect of dsDNA on the Assembly Pathway and Mechanical Strength of SV40 VP1 Virus-like Particles. Biophys. J. 115, 1656–1665 (2018).

39. S. F. J. Wickham, A. Auer, J. Min, N. Ponnuswamy, J. B. Woehrstein, F. Schueder, M. T. Strauss, J. Schnitzbauer, B. Nathwani, Z. Zhao, S. D. Perrault, J. Hahn, S. Lee, M. M. Bastings, S. W. Helmig, A. L. Kodal, P. Yin, R. Jungmann, W. M. Shih, Complex multicomponent patterns rendered on a 3D DNA-barrel pegboard. Nat. Commun. 11, 5768 (2020).

40. C. F. Bohren, D. R. Huffman, Absorption and Scattering of Light by Small Particles. doi: 10.1002/9783527618156 (1983).

41. E. van der Pol, F. A. W. Coumans, A. Sturk, R. Nieuwland, T. G. van Leeuwen, Refractive Index Determination of Nanoparticles in Suspension Using Nanoparticle Tracking Analysis. Nano Lett. 14, 6195–6201 (2014).

42. Y. Pang, H. Song, W. Cheng, Using optical trap to measure the refractive index of a single animal virus in culture fluid with high precision. Biomed. Opt. Express 7, 1672–1689 (2016).

43. L. Ma, S. Zhu, Y. Tian, W. Zhang, S. Wang, C. Chen, L. Wu, X. Yan, Label-Free Analysis of Single Viruses with a Resolution Comparable to That of Electron Microscopy and the Throughput of Flow Cytometry. Angew. Chem. Int. Ed. 128, 10395–10399 (2016).

44. J. Vörös, The Density and Refractive Index of Adsorbing Protein Layers. Biophys. J. 87, 553–561 (2004).

45. M. Röding, H. Deschout, K. Braeckmans, M. Rudemo, Measuring absolute number concentrations of nanoparticles using single-particle tracking. Phys. Rev. E 84, 031920 (2011).

46. Y. Xiong, X. Zhang, L. Hu, A method for tracking the Brownian motion to estimate the size distribution of submicron particles in seawater. Limnol. Oceanogr.: Methods 20, 373–386 (2022).

47. B. Hua, K. Y. Han, R. Zhou, H. Kim, X. Shi, S. C. Abeysirigunawardena, A. Jain, D. Singh, V. Aggarwal, S. A. Woodson, T. Ha, An improved surface passivation method for single-molecule studies. Nat. Methods 11, 1233–1236 (2014).

48. K. Takamura, H. Fischer, N. R. Morrow, Physical properties of aqueous glycerol solutions. J. Pet. Sci. Eng. 98, 50–60 (2012).

49. T. Galpin, R. T. Chartier, N. Levergood, M. E. Greenslade, Refractive index retrievals for polystyrene latex spheres in the spectral range 220–420 nm. Aerosol Sci. Technol. 51, 1158–1167 (2017).

50. M. A. Odete, F. C. Cheong, A. Winters, J. J. Elliott, L. A. Philips, D. G. Grier, The role of the medium in the effective-sphere interpretation of holographic particle characterization data. Soft Matter 16, 891–898 (2020).

